# Draft Genome Assemblies and Annotations of *Agrypnia vestita* Walker, and *Hesperophylax magnus* Banks Reveal Substantial Repetitive Element Expansion in Tube Case-making Caddisflies (Insecta: Trichoptera)

**DOI:** 10.1101/2020.11.16.381806

**Authors:** Lindsey K. Olsen, Jacqueline Heckenhauer, John S. Sproul, Rebecca B. Dikow, Vanessa L. Gonzalez, Matthew P. Kweskin, Adam M. Taylor, Seth B. Wilson, Russell J. Stewart, Xin Zhou, Ralph Holzenthal, Steffen U. Pauls, Paul B. Frandsen

## Abstract

Trichoptera (caddisflies) play an essential role in freshwater ecosystems; for instance, larvae process organic material from the water and are food for a variety of predators. Knowledge on the genomic diversity of caddisflies can facilitate comparative and phylogenetic studies thereby allowing scientists to better understand the evolutionary history of caddisflies. While Trichoptera are the most diverse aquatic insect order, they remain poorly represented in terms of genomic resources. To date, all long-read based genomes have been sequenced from individuals in the retreat-making suborder, Annulipalpia, leaving ∼275 Ma of evolution without high-quality genomic resources. Here, we report the first long-read based *de novo* genome assemblies of two tube case-making Trichoptera from the suborder Integripalpia, *Agrypnia vestita* Walker and *Hesperophylax magnus* Banks. We find that these tube case-making caddisflies have genome sizes that are at least three-fold larger than those of currently sequenced annulipalpian genomes and that this pattern is at least partly driven by major expansion of repetitive elements. In *H. magnus*, long interspersed nuclear elements (LINEs) alone exceed the entire genome size of some annulipalpian counterparts suggesting that caddisflies have high potential as a model for understanding genome size evolution in diverse insect lineages.

**Significance:** There is a lack of genomic resources for aquatic insects. So far, only three high-quality genomes have been assembled, all from individuals in the retreat-making suborder Annulipalpia. In this article, we report the first high-quality genomes of two case-making species from the suborder Integripalpia, which are essential for studying genomic diversity across this ecologically diverse insect order. Our research reveals larger genome sizes in the tube case-makers (suborder Integripalpia, infraorder Phryganides), accompanied by a disproportionate increase of repetitive DNA. This suggests that genome size is at least partly driven by a major expansion of repetitive elements. Our work shows that caddisflies have high potential as a model for understanding how genomic diversity might be linked to functional diversification and forms the basis for detailed studies on genome size evolution in caddisflies.

**Data deposition:** This project has been deposited at NCBI under the Bioproject ID: PRJNA668166

## Introduction

With 16,544 extant species (Morse 2020), caddisflies (Insecta: Trichoptera) are the most diverse of the primary aquatic insect orders, comprising more species than the other four (Odonata, Ephemeroptera, Plecoptera, and Megaloptera) combined (Dijkstra et al., 2014). This diverse group of insects has successfully colonised all types of freshwater (and even intertidal) habitats across all continents north of Antarctica. Within these freshwater ecosystems caddisflies play important roles, including nutrient cycling and energy flow, and stabilizing the waterbed. They also act as biological indicators of water quality (Morse et al., 2019). Trichoptera is divided into two suborders, Annulipalpia and Integripalpia, both of which produce silk in modified labial glands (Thomas et al., 2020b). Annulipalpians use silk to construct small homes and capture nets that are fixed to the substrate, whereas most integripalpians’ use silk to connect material into portable tube cases offering protection, camouflage, and even aiding in respiration (Fig. 1) (Wiggins, 2004). This innovation in extended phenotype has potentially facilitated their radiation across a multitude of different environments including streams, lakes, ponds, and even marine environments.

**Figure 1.**
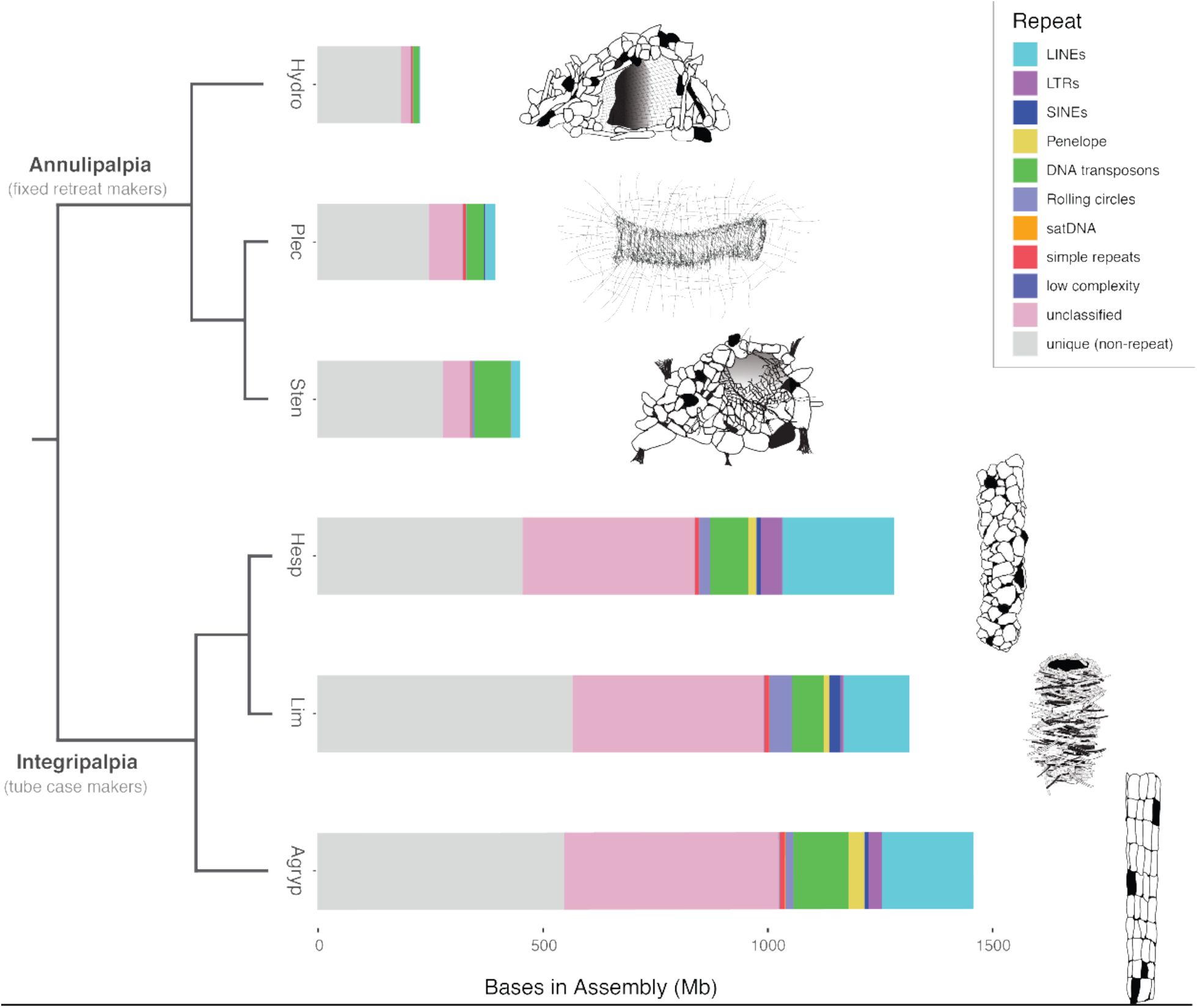
Assembly length and repetitive DNA content in Trichoptera suborders. Comparison of assembly length and repetitive DNA content among genome assemblies for three annulipalipan (*Hydropsyche tenuis, Plectrocnemia conspersa*, and *Stenopsyche tienmushanensis*) and three integripalpian (*Hesperophylax magnus, Limnephilus lunatus*, and *Agrypnia vestita*) species. The total length of bars indicates assembly size and colored segments within bars indicate the fraction of the assembly belonging to major repeat categories identified by RepeatModeler2 and annotated by RepeatMasker. Artwork to the right of plots shows examples of fixed retreats built by annulipalpians compared to integripalpian tube cases. Each Illustration is derived from a member of the same genus as the genome assemblies.

Relative to their diversity, most insect groups remain poorly represented in existing genomic resources—a trend which is particularly pronounced in aquatic insects (Hotaling et al., 2020). Yet insects commonly show dynamic genome evolution within groups, including major variation in genome size that is often linked to expansion and loss of repetitive DNA (Lower et al., 2017; Petersen et al., 2019; Pflug et al., 2020). Insect diversity offers a vast supply of potential model systems for understanding how genomes evolve, especially as advancing sequencing technology enables more cost-effective, high-quality genome assemblies in any model system. Currently, there are three long-read based draft Trichoptera genome assemblies (Luo et al., 2018; Heckenhauer et al., 2019; Table 1). However, all of these were generated for species within the suborder Annulipalpia, leaving approximately 10,453 Integripalpian species (Morse 2020) and 275 million years of evolutionary history poorly represented by genomic resources (Thomas et al., 2020b). In addition, a lack of genetic resources in the large case-making radiation within caddisflies prevents research into the genomic basis of the fascinating evolutionary history and ecological diversification of this diverse and important group of caddisflies. Here, we report the first long-read based *de novo* genome assemblies and annotations of two tube-case making intergripalpian caddisflies, *Agrypnia vestita* Walker and *Hesperophylax magnus* Banks. Their estimated genome sizes are more than 3-fold larger than previously sequenced annulipalpian caddisflies. We show that this is, at least partly, due to a large expansion of repeat content in the case-making caddisflies compared to retreat-making caddisflies.

**Table 1.** Comparison of Genome Assemblies against Previously Published Caddisfly Genomes.

| Species | Accession | Suborder | Sequencing Platform Coverage | Assembly Length(bp) | Contig N50 (kbp) |
| --- | --- | --- | --- | --- | --- |
| *Agrpynia vestita* (present study) | JADDOH000000000 | Integripalpia | PacBio+Illumina (17.86x + 87.96x) | 1,352,945,503 | 111.8 |
| *Hesperophyla x magnus* (present study) | JADDOG000000000 | Integripalpia | Nanopore+Illumina (26.38x +49.30x) | 1,233,588,871 | 768.2 |
| *Hydropsyche tenuis* (Heckenhauer et al., 2019) | GCA_009617725.1 | Annulipalpia | Nanopore+Illumina (16.5x +167.6x) | 229,663,394 | 2190.1 |
| *Plectrocnemia conspersa* (Heckenhauer et al., 2019) | GCA_009617715.1 | Annulipalpia | Nanopore+Illumina (17.1x + 82.9x) | 396,695,105 | 869.0 |
| *Stenopsyche tienmushanensis* (Luo et al., 2018) | GCA_008973525.1 | Annulipalpia | PacBio+Illumina (153x + 150x) | 451,494,475 | 1296.9 |
| *Limnephilus lunatus* (Thomas et al., 2020a) | Llun_2.0 | Integripalpia | Illumina (80.1x) | 1,269,180,260 | 24.2 |
| *Glossosoma conforme* (Weigand et al., 2018) | GCA_003347265.1 | Annulipalpia | Illumina (53x) | 604,293,666 | 14.2 |
| *Sericostoma* sp. (Weigand et al., 2017) | GCA_003003475.1 | Integripalpia | Illumina (43x) | 1,015,727,762 | 2.1 |

<sup>a</sup> Present = complete + fragmented
<sup>b</sup> N<sub>Insecta</sub> = 1367

## Materials and Methods

### Sequencing and assembly

We collected individuals of both species in the wild, *A. vestita* as an adult and *H. magnus* as a pupa. Following extraction, we sequenced genomic DNA from *A. vestita* on an HiSeq 2500 lane and on 23 PacBio sequel SMRT cells. We sequenced genomic DNA from *H. magnus* using two Illumina NovaSeq and four Oxford Nanopore FLO-MIN 106 flow cells. Further details are provided in Supplementary Note 1, Supplementary Material online. For both data sets, we conducted a *de novo* hybrid assembly using MaSuRCA v.3.1.1 (Zimin et al., 2013, 2017). MaSuRCA aligns high-fidelity short reads to more noisy long reads to generate “megareads”, which are then assembled in CABOG (Miller et al., 2008), an overlap-layout-consensus assembler. In the config file for each run, we specified an insert size of 500 bps for the Illumina paired-end reads with a standard deviation of 50 and a Jellyfish hash size of 100,000,000,000. All other parameters were left as defaults. We screened genome assemblies for potential contaminants with BlobTools v1.0 (Laetsch and Blaxter, 2017; Supplementary Note 2, Supplementary Material online). Contigs consisting of contaminant DNA were subsequently removed from the final assemblies. We assessed genome quality and completeness with BUSCO v4.1.1 (Seppey et al., 2019; Supplemental Note 3, Supplementary Material online) with the OrthoDB v.10 Insecta and Endopterygota gene sets (Kriventseva et al., 2019) and generated genome statistics using the assembly_stats script (Trizna, 2020; Supplemental Table 1, Supplementary Material online for full output). We conducted genome profiling (estimation of major genome characteristics such as size, heterozygosity, and repetitiveness) on the short-read sequence data with GenomeScope 2.0 (Ranallo-Benavidez et al., 2020) as described in in Supplementary Note 4, Supplementary Material online.

### Repeat and Gene Annotation

We conducted comparative analysis of repetitive elements for the genomes generated in this study, the three available long-read annulipalpian genomes, and *Limnephilus lunatus*, the integripalpian with the highest quality short-read genome assembly. We identified and classified repetitive elements *de novo* and generated a library of consensus sequences using RepeatModeler 2.0 (Flynn et al., 2019). We then annotated repeats in the assembly with RepeatMasker 4.1.0 (Smit and Hubley, 2008–2015) using the custom repeat library generated in the previous step. We conducted an orthogonal analysis of repeat dynamics using a reference-free approach by normalizing subsampled Illumina data for each sample using RepeatProfiler (Negm et al., 2020) and then analyzing normalized data sets for repeat content in RepeatExplorer2 (Novák et al., 2013), with more details provided in Supplementary Note 5, Supplementary Material online.

To generate evidence for gene annotation, we aligned previously sequenced transcriptomes to each genome using BLAST-like Alignment Tool v3.6 (BLAT, Kent, 2002). We aligned the transcriptome of the closely related *Phryganea grandis* from the 1KITE project (111126_I883_FCD0GUKACXX_L7_INShauTBBRAAPEI-22 http://www.1kite.org/) to *A. vestita*, and we aligned the *Hesperophylax* transcriptome from (Wang et al., 2015) to *H. magnus*. We generated *ab initio* gene predictions using AUGUSTUS v3.3 (Stanke et al., 2008) with hints generated from RepeatMasker 4.1.0 and BLAT v3.6 and by supplying the retraining parameters obtained from the BUSCO analysis (Supplementary Note 6, Supplementary Material online). Following annotation, we removed genes from our annotation that did not generate significant BLAST hits or lacked transcript evidence. Lastly, functional annotations were identified using Blast2GO (Götz et al., 2008).

## Results and Discussion

### Assembly

Here, we generated the first genome assemblies based on long-read sequencing from the species diverse caddisfly suborder, Integripalpia. They provide important genome resources and fill a gap in evolutionary history of more than 275 million years (Thomas et al., 2020b). The *A. vestita* genome was sequenced using ∼88x Illumina sequence coverage and ∼18x PacBio read coverage. After contaminated contigs were removed, the resulting assembly contained 25,541 contigs, a contig N50 of 111,757 bp, GC content of 33.77%, and a total length of 1,352,945,503 bp. BUSCO analysis identified 94.4% (91.4 % complete, 3.0% fragmented) of the Insecta gene set in the assembly (see Supplemental Note 3, Supplementary Material online for further details). The *H. magnus* genome was sequenced with ∼49x Illumina sequence coverage and ∼26x Oxford Nanopore sequence coverage. The resulting assembly has 6,877 contigs, a contig N50 of 768,217bp, GC content of 34.36%, and a total length of 1,275,967,528 bp. We identified 95.9% (95.2% complete, 0.7% fragmented) of the Insecta BUSCO gene set in the final assembly. Although the sequencing and assembly techniques were similar to those used in previous efforts to sequence and assemble high quality reference genomes in Trichoptera (Table 1, Heckenhauer et al., 2019; Luo et al., 2018), the contiguity of these genomes was lower. This is likely to have been caused by large genome size and the proliferation of repetitive DNA, which represents one of the primary barriers to genome assembly. However, despite these challenges, both genome assemblies represent a substantial improvement in contiguity to previous assemblies of integripalpian caddisflies generated from short read data alone. For example, at the time of writing, the highest quality integripalpian genome assembly on GenBank is *Limnephilus lunatus*, which was assembled from short read data and has a contig N50 of 24.2 kb, giving further evidence to the difficulty of assembling large, repetitive caddisfly genomes.

### Annotation and Repeat Analysis

We also report the functional annotations of *H. magnus* and *A. vestita*. Of 59,600 proteins predicted by AUGUSTUS for *A. vestita*, 21,637 were verified by BLAST and/or transcript evidence (and maintained in the final annotation), 14,096 were mapped to GO terms, and 5,362 were functionally annotated in BLAST2GO. Of 38,490 proteins predicted by AUGUSTUS for *H. magnus*, 16,791 were verified by BLAST and/or transcript evidence (and maintained in the final annotation), 10,605 were mapped to GO terms, and 5,362 were functionally annotated in BLAST2GO. Top GO annotations include cellular process (*A. vestita* 3395, *H. magnus* 2366), metabolic process (*A. vestita* 2897, *H. magnus* 2047), binding (*A. vestita* 3425, *H. magnus* 2187), and catalytic activity (*A. vestita 3243, H. magnus* 2395) (Supplemental Figure 6 and 7, Supplementary Material online).

The results of genome assembly repeat annotation, genome profiling, and *de novo* repeat assembly with RepeatExplorer2 all showed a disproportionate increase of repetitive DNA in integripalpian genomes compared to annulipalpians for those species sampled (Fig. 1, Supplementary Note 5, Supplementary Figs. 3 and 4, Supplementary Material online). In the integripalpian species, unclassified repeats alone make up an average of >400 million bases, which exceeds the average estimated genome size of all three annulipalpians analyzed. After unclassified repeats, long interspersed nuclear elements (LINEs) are the most abundant repeat category showing a disproportionate increase. LINEs comprise an average of >200 million bases in integripalpians, and show a ∼4-fold average increase in genome proportion (avg. genome proportion = 15.1%) compared to the annulipalpians (avg. genome proportion = 3.8%). *Hesperophylax* has more bases annotated as LINEs (∼249 million) than the size of the entire *Hydropsyche* genome assembly. Rolling-circles and long terminal repeats (LTRs) also show disproportionate increase in integripalpians (∼3.5-fold and 18-fold increases in genome proportion, respectively), however both categories make up a much smaller fraction of integripalipan genomes (<2.5% on average). DNA transposons are abundant in all integripalpian genomes we studied (average of 92 million bases annotated), however their genomic proportion decreased relative to annulipalipans in which DNA transposons were the most abundant classifiable repeat category (11.3% avg. genome proportion in annulipalipans vs 6.8% in integripalpians).

The high abundance of unclassified repeats we observed in the integripalpian genomes is not surprising given that Trichoptera repeats are poorly represented in repeat databases. Unclassified repeats may also represent the remnants of ancient transposable element expansions, which are particularly difficult to annotate (Hoen et al., 2015). This explanation of old repeat expansions accounting for much of the unclassified repeats is consistent with results of clustering analysis in RepeatExplorer2 which shows many unannotated superclusters that make up small fractions of the genome (Supplemental Fig. 3, Supplementary Material online). We do not observe large unannotated superclusters that would indicate failed annotation of abundant, recently active repeats. Given the apparent suborder-specific increase in unclassified repeats, we hypothesize that ancient transposable element activity in the ancestor of integripalpians contributed to the larger genome sizes we observe, however denser sampling of genomes across Trichoptera suborders is needed to address this hypothesis..

Given the major variation in genome size and repeat abundance, our findings suggest Trichoptera has high potential as a model for gaining insights into genome evolution in diverse insect lineages. Future investigation on the role of LINEs in genome diversification is of particular interest given our findings. We present preliminary evidence that LINEs show suborder-specific expansions, albeit with very limited taxon sampling. LINEs (especially L1) play major roles in genome stability, cancer, and aging (De Cecco et al., 2019; Van Meter et al., 2014). In many groups LINEs are hypothesized to play important evolutionary roles, including roles in rapid genome evolution though their own movement (Kordiš et al., 2006; Suh et al., 2015; Warren et al., 2008), and by facilitating expansion of other repeat classes (Grandi and An, 2013; Sproul et al., 2020). It is possible that these elements have been important drivers in the expansion of integripalpian genomes. The high-quality genome assemblies and repeat libraries we present here provide a starting point for investigating the role of repeats in genome evolution across caddisfly lineages. In addition, we close a large evolutionary gap in genomic resources within a large, ecologically diverse clade in which additional genome sequencing can enable new insights as to the genomic basis of adaptation and diversification within freshwater environments.

## Acknowledgments

This material is based upon work supported by the Global Genome Initiative under Grant No. (GGI-Exploratory-2016-049). J.H was supported by LOEWE-Centre for Translational Biodiversity Genomics funded by the Hessen State Ministry of Higher Education, Research and the Arts (HMWK). JSS was supported by a NSF Postdoctoral Research Fellowship in Biology (DBI-1811930). Part of the computation was performed on the Smithsonian High-Performance Computing Cluster (SI/HPC: doi.org/10.25572/SIHPC). L.K.O. was supported by a BYU College of Life Sciences Undergraduate Research Award.

## Authors’ Contributions

*Conceptualization:* LKO, JH, VLG, MPK, RBD, SUP, JSS, PBF

*Methodology:* LKO, JH, JSS, VLG, MPK, RBD, SUP, PBF

*Software:* AMT

*Validation:* LKO, JSS, PBF

*Formal analysis:* JH, JSS, LKO, RBD, PBF

*Investigation:* VLG, AMT, SW, PBF

*Resources:* VLG, RH, RJS, PBF

*Writing-original draft preparation:* LKO,JH, JSS, LKO, PBF

*Writing-review and editing:* LKO, JH, MPK, RBD, JSS, SUP, AMT, RJS, RH, XZ, PBF

*Visualization:* JSS, RH

*Supervision:* PBF

*Project administration:* LKO, PBF

*Funding acquisition:* LKO, VLG, MPK, RBD, SUP, RJS, XZ, PBF

## Notes

### Competing Interest Statement

The authors have declared no competing interest.

