## Supplemental_Material for "Draft Genome Assemblies and Annotations of *Agrypnia vestita* Walker, and *Hesperophylax magnus* Banks Reveal Substantial Repetitive Element Expansion in Tube Case-making Caddisflies (Insecta: Trichoptera)"

Supplementary Note 1: **DNA extraction, library preparation, and sequencing**

*Agrypnia vestita*

An adult *A. vestita* was collected in Ramsey County, Roseville, Minnesota USA (45.0325°N, 93.17793°W) and immediately flash-frozen. High-molecular-weight genomic DNA was extracted using the Qiagen MagAttract HMW DNA Kit from the snap-frozen head and thorax according to the manufacturer's directions, performed at the Laboratories of Analytical Biology at the Smithsonian Institution’s National Museum of Natural History (NMNH). HMW Genomic DNA was quantified by fluorometry (Qubit, Thermo Fisher Scientific, Waltham, U.S.A.) and assessed for size by 1.0% (w/v) agarose gel using Pulse Field Electrophoresis (PFGE) agarose gel electrophoresis (Chef Mapper XA, Bio-Rad Laboratories, Inc).

Single Molecule Real Time (SMRT) bell libraries were prepared according to the "20 kb Template Preparation Using BluePippin Size-Selection System" as recommended by Pacific Biosciences (https://www.pacb.com/wp-content/uploads/2015/09/Procedure-Checklist-20-kb-Template-Preparation-Using-BluePippin-Size-Selection.pdf, Palo Alto, U.S.A). After damage-repair the libraries were size-selected on a BluePippin system (0.75% (w/v) agarose gel cassette, dye-free, S1 marker, high pass 20kb protocol) to remove library fragments smaller than 8 kb. Then libraries were recovered by PB AMPure beads, quantified by the high sensitivity fluorometric assay (Qubit, Thermo Fisher Scientific, Waltham, U.S.A.) and quality assessed using the genomic assay on the TapeStation (Agilent, Waldbronn, Germany). SMRT bell templates were bound to P6 polymerase using the DNA polymerase binding kit P6 v2 primers. Polymerase-template complexes were bound to magnetic beads using the Magbead Binding Kit and sequencing was carried out on the PacBio RS II (23 cells) sequencer with movie lengths of 360 min.

Illumina library preparation and sequencing was performed by Novogene using Nextera library prep followed by 2x150 paired-end sequencing on an Illumina HiSeq 2500.

*Hesperophylax magnus*

We collected a *H. magnus* pupa in Salt Lake County, Utah, USA in the Red Butte Creek (40.774273°N 111.817617°W). We extracted genomic DNA using the Agilent DNA extraction kit and prepared libraries and sequenced them using both paired-end sequencing on an Illumina NovaSeq and on four Oxford Nanopore FLO-MIN 107 flow cells using the MinION portable DNA sequencer and the LSK-109 ligation library kit.

Supplementary Note 2: **Contamination-screening using BlobTools**

The final genome assemblies were screened for potential contaminations with taxon annotated GC-coverage (TAGC) plots using BlobTools v1.0. For this purpose, all Illumina reads were mapped against the final genome assemblies using BWA-MEM v0.7.17-r1188 (Li, 2013). Taxonomic assignment for BlobTools was done with blastn using -task megablast and -e-value 1e-25. Contigs which had a blast hit with Chordata (179 in *H. magnus*, 300 in *A. vestita*) and Cnidaria (1,054 in *A. vestita*) were filtered out (Supplemental Figure 1 and 2). The total length of the contamination filtered assemblies was 1,233,588,871 bp and 1,352,945,503 bp for the *H. magnus* and *A. vestita* assembly respectively.

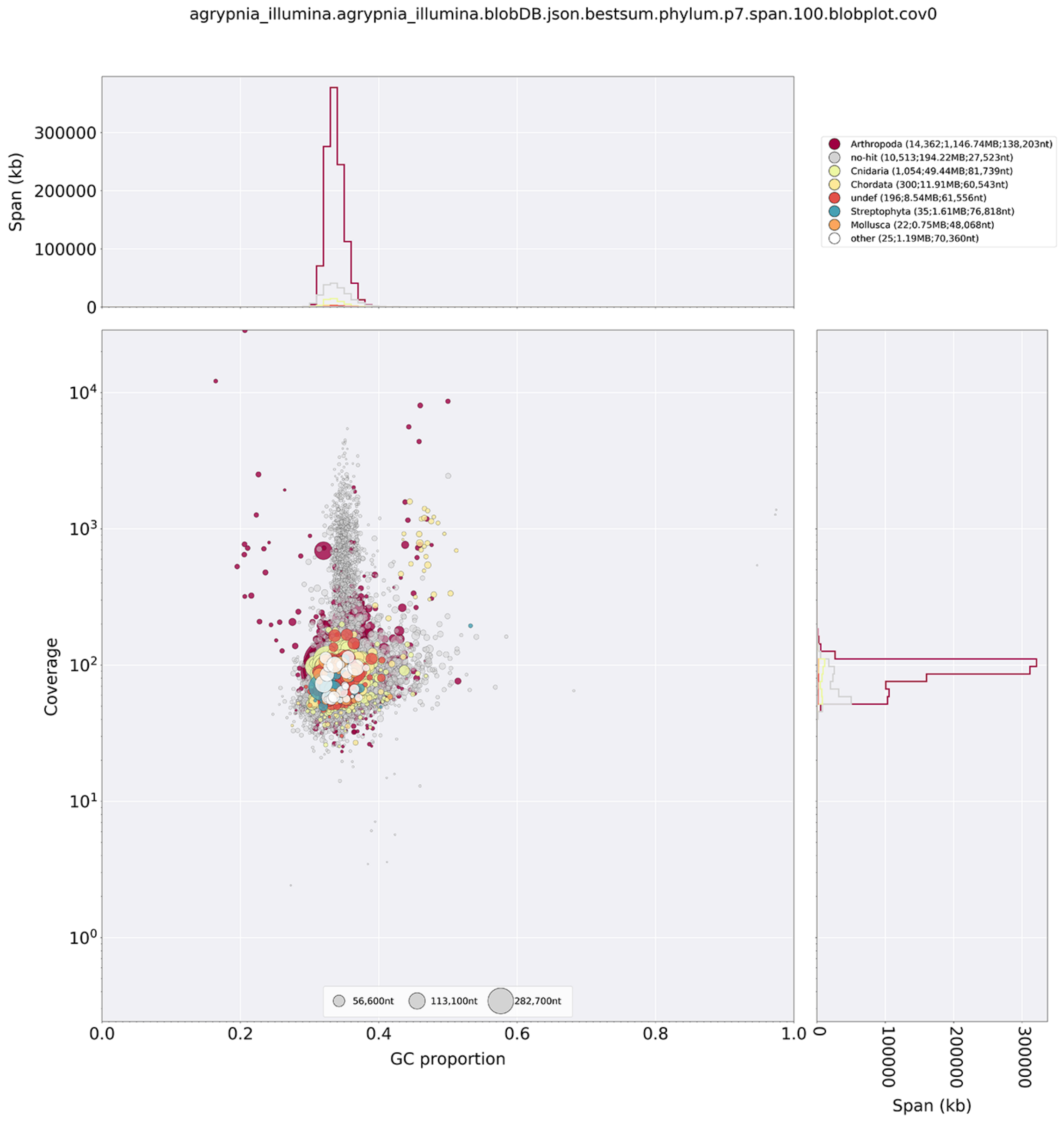


**Supplemental Figure 1.** Taxon-annotated GC-coverage (TAGC) plots for pre-filtered *A. vestita* genome assemblies. Circles indicate contigs and the color indicates the best match to taxon annotation. The upper and right hand panel show the total span of contigs (kb) given GC proportion.

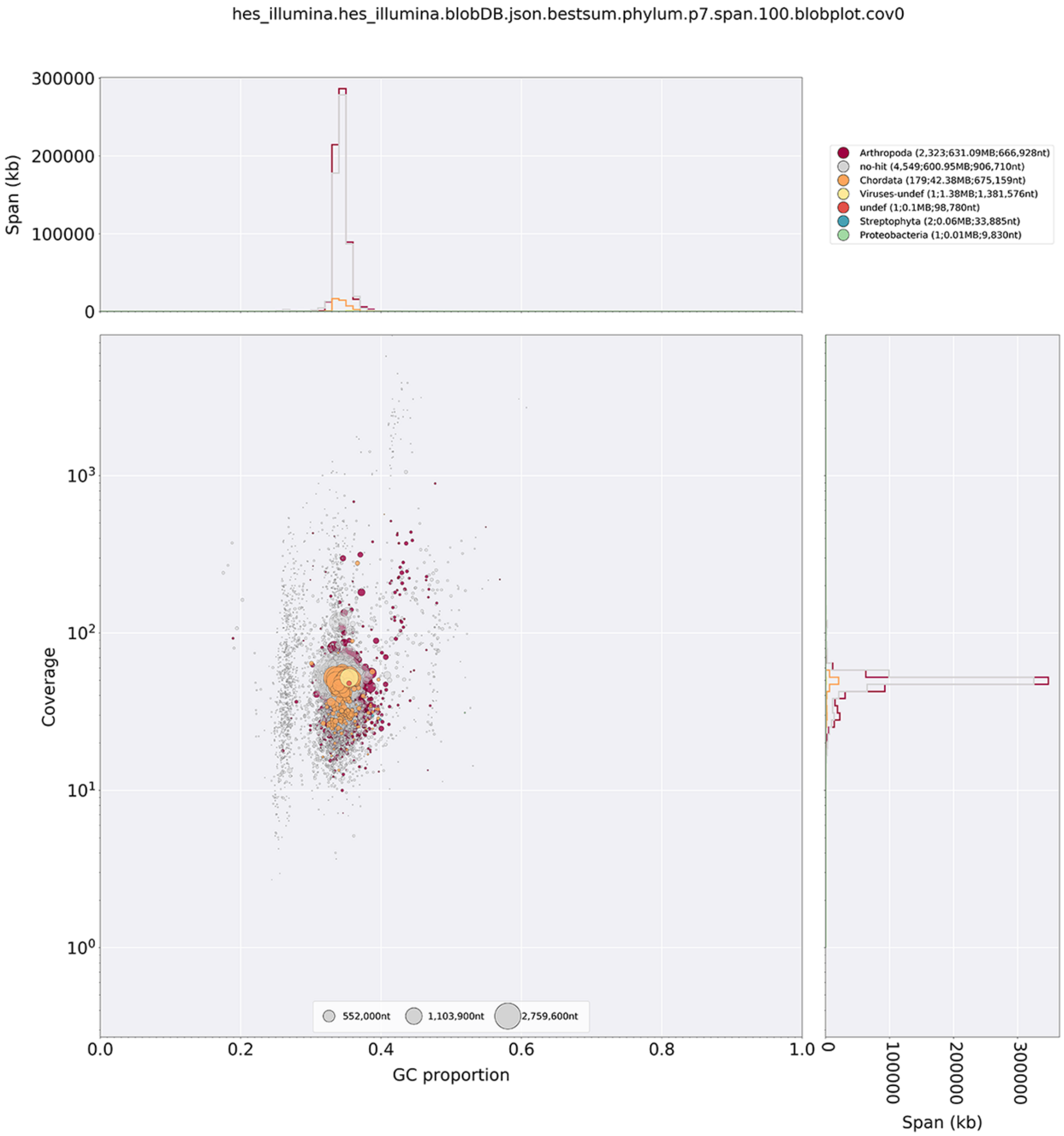


**Supplemental Figure 2.** Taxon-annotated GC-coverage (TAGC) plots for pre-filtered *H. magnus* genome assemblies*.* Circles indicate contigs and the color indicates the best match to taxon annotation. The upper and right hand panel show the total span of contigs (kb) given GC proportion.

Supplementary Note 3: **BUSCO analysis**

BUSCO comparisons between *Agrypnia vestita* and *Hesperophylax magnus* filtered assemblies and previous Trichoptera assemblies were done using BUSCO v4.1.1 (Seppey et al., 2019) and the insecta_odb10 dataset (https://busco-data.ezlab.org/v4/data/lineages/insecta_odb10.2020-08-05.tar.gz) with options --long -m genome and --offline. The results are summarized in table 1. BUSCO v.4.1.1 was also run using the endopterygota_odb10 dataset (https://busco-data.ezlab.org/v4/data/lineages/endopterygota_odb10.2020-08-05.tar.gz). In total, BUSCO detected 93.3% of the Endopterygota core gene collection in the predicted proteins of *A. vestita* (complete: 88.9%, fragmented: 4.4%). For *H. magnus* 94.9% of the genes were detected (complete: 93.6%, fragmented: 1.6%).

**Table 1.** Assembly statistics for filtered assemblies. Assembly stats were calculated using the assembly_stats.py function (Mike Trizna, 2020)

| **Contig Stats** | ***A. vestita*** | ***H. magnus*** |
|---|---|---|
| L10 | 267 | 26 |
| L20 | 711 | 79 |
| L30 | 1319 | 158 |
| L40 | 2134 | 262 |
| L50 | 3196 | 401 |
| N10 | 368769 | 3008714 |
| N20 | 359634 | 1845881 |
| N30 | 190978 | 1372687 |
| N40 | 14882 | 1038400 |
| N50 | 111757 | 768217 |
| GC content | 33.77 | 34.36 |
| Longest | 1130755 | 11038151 |
| Mean | 52971.52 | 179378.93 |
| Median | 25612.0 | 34916.0 |
| Sequence count | 25541 | 6877 |
| Shortest | 1057 | 1353 |
| Total bps | 1352945503 | 1233588871 |

Supplementary Note 4 **Genome profiling based on a k-mer distribution-based method**

For genome scope profiling, Illumina reads generated in this study, as well as previously published data was used. Illumina reads of *Limnephilus lunatus* (SRR947083) and

*Stenopsyche tienmushanensis* (SRR7062469) were downloaded from the European Nucleotide Archive (https://www.ebi.ac.uk/ena) and raw reads of *Hydropsyche tenuis* (SRS5312808) and Plectrocnemia conspersa (SRS5312807) were obtained from SRA (https://www.ncbi.nlm.nih.gov/sra). Raw reads were contamination filtered using kraken2 v 2.0.8-beta with the default database. Before running GenomeScope 2.0 (Ranallo-Benavidez et al., 2020), k-mers were counted with JELLYFISH v2.2.10 (Marçais and Kingsford, 2011) using jellyfish count -C -s 25556999998 -F 3 and a k-mer length of 21 (-m 21) as recommended for most genomes by the authors of GenomeScope with the contamination filtered Illumina reads. A histogram of k-mer frequencies was produced with jellyfish histo. GenomeScope 2.0 was run with the exported k-mer count histogram within the online web tool (http://qb.cshl.edu/genomescope/genomescope2.0/) using the following parameters:

Kmer length = 21, Max kmer coverage = 10000. Resulting GenomeScope2

profiles are available online:

**Table 2.** GenomeScope2 Results.

| **Species** | **Genome size (bp)** | **Unique (%)** | **Link to Genomescope2 profile** |
|---|---|---|---|
| *Agrypnia vestita* | 931,685,836 | 68.3 | http://qb.cshl.edu/genomescope/genomescope2.0/analysis.php?code=sbLuS8k74GWviwYJBMcC |
| *Hesperophylax magnus* | 1,060,786,686 | 57.4 | http://qb.cshl.edu/genomescope/genomescope2.0/analysis.php?code=Mt4Qb8mdRzEafda5zh0x |
| *Limnephilus lunatus* | 1,080,975,116 | 53.4 | http://qb.cshl.edu/genomescope/genomescope2.0/analysis.php?code=MxZkVb4nqGegS0lolPsC |
| *Stenopsyche tienmushanensis* | 389,476,263 | 85.1 | http://qb.cshl.edu/genomescope/genomescope2.0/analysis.php?code=w6vjdQTYYElr2cTsp3zK |
| *Plectrocnemia conspersa* | 316,263,089 | 98.1 | http://qb.cshl.edu/genomescope/genomescope2.0/analysis.php?code=Xuq6HXkomG412S7ksIWC |
| *Hydropsyche tenuis* | 222,816,390 | 92.6 | http://qb.cshl.edu/genomescope/genomescope2.0/analysis.php?code=swsSPZYIOejH8dfenSFm |

Supplementary Note 5: **Repeat Analysis**

We further explored repeat dynamics with a reference-free approach using RepeatExplorer2 (Novák et al., 2013) and TAREAN (Novák et al., 2017). This orthogonal approach which estimates repeat abundance directly from short-read data can show improved quantification of repeats prone to underrepresentation in genome assemblies, such as large blocks of satellite DNA which may be present in some species. Prior to analysis with RepeatExplorer2 we normalized input reads across samples by mapping contamination filtered Illumina reads to the Endopterygota BUSCO gene set using RepeatProfiler (Negm et al., 2020), and used average coverage values of BUSCO genes to calculate the number of reads required for 0.5X coverage of BUSCO genes per sample. To avoid bias introduced by BUSCO genes with 0 coverage genes as well as those with unexpectedly high coverage (e.g., due to unexpected mapping of repetitive sequences), we calculated 0.5x coverage using only the middle 70% of the BUSCO gene coverage values when genes were ordered by maximum coverage depth. We downsampled reads using seqtk (Li, 2020). We summarized RepeatExplorer2 output by plotting the abundance of major repeat categories present in top clusters in R (Team, R. Core)

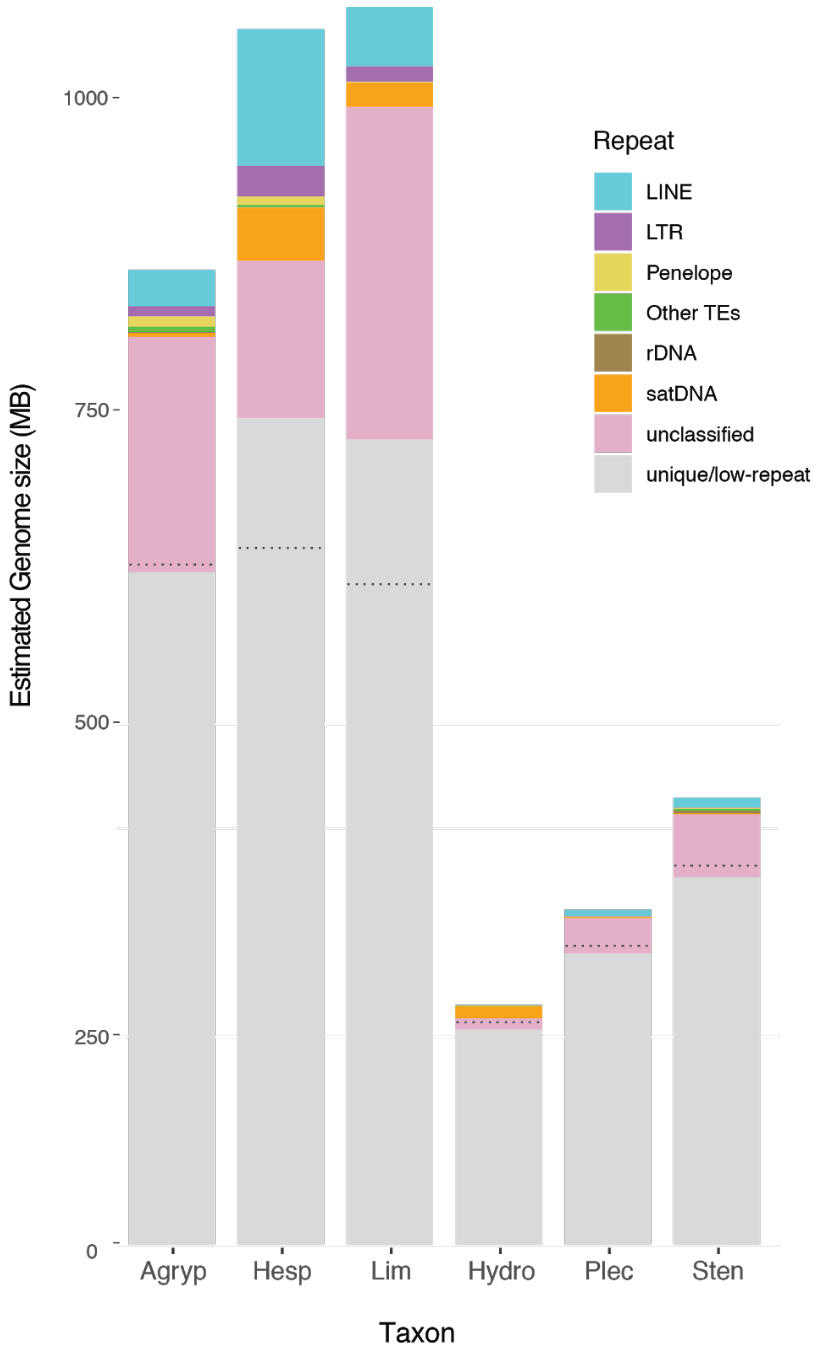


**Supplemental Figure 3:** Comparison of genome size and repetitive DNA content in Trichoptera suborders estimated *de novo* from short-read sequences. Estimates include three integripalpian (Hesperophylax, Limnephilus, and Agrypnia) and three annulipalipans (Hydropsyche, Plectrocnemia, and Stenopsyche). The height of bars indicates genome size estimates obtained using GenomeScope and the colored segments within bars indicate the genomic proportion of major repeat categories identified by clustering analysis in RepeatExplorer2. The dotted line indicates the threshold of unique vs repetitive DNA sequences estimated by GenomeScope (kmer size = 21).

–

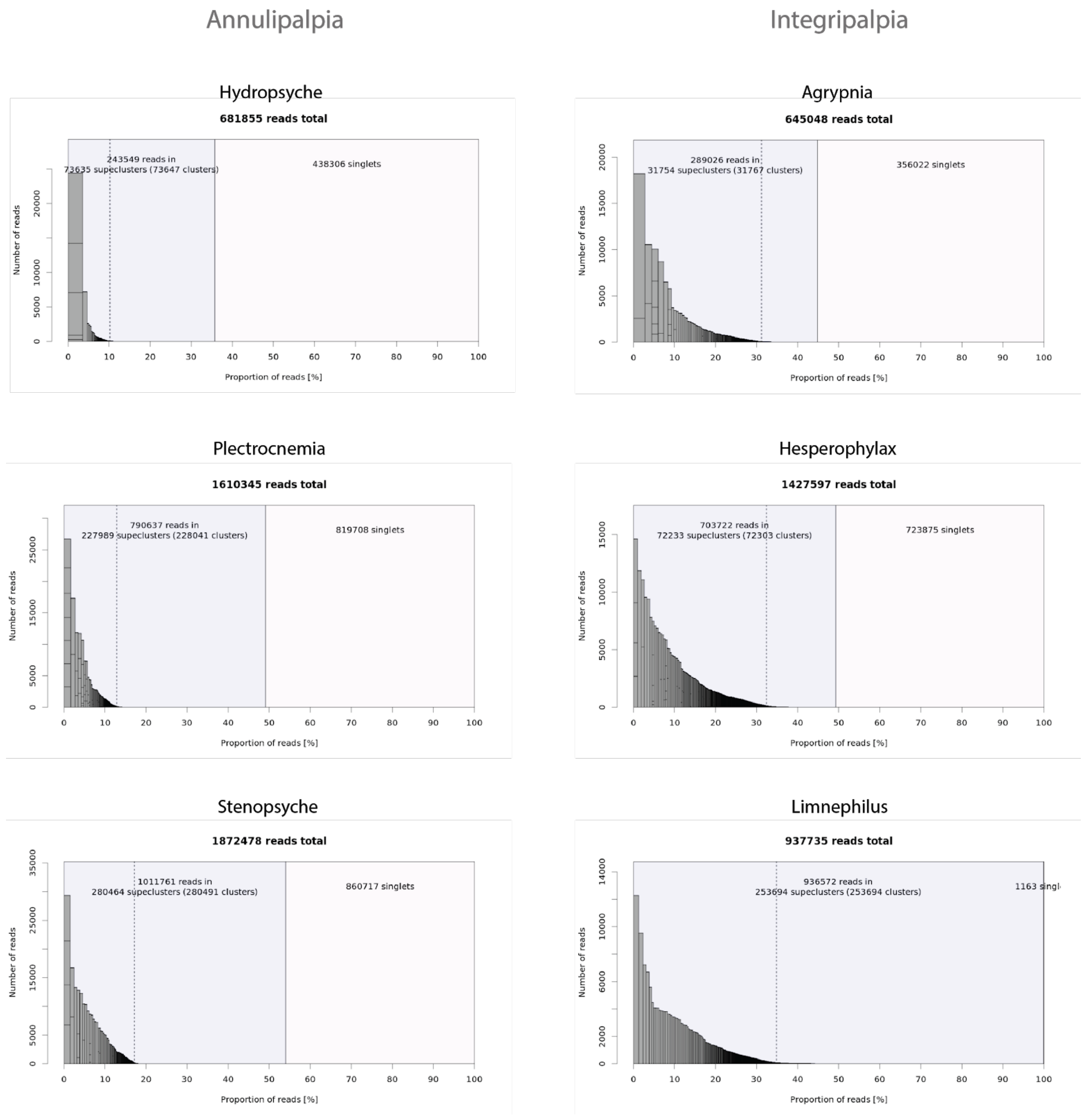


**Supplemental Figure 4:** Summary of RepeatExplorer2 clustering analysis. Gray bars represent superclusters of repetitive sequences with their heights and widths corresponding to the number of reads present in that supercluster (see y-axis). The genomic proportion of each supercluster is shown on the x-axis with top-clusters shown to the left of the dotted line. The fraction of repetitive clusters vs singlet reads is shown by the purple and pink boxes, respectively.

Supplementary Note 6: **Genome Annotation**

Annotations for *Agrypnia vestita* and *Hesperophylax magnus* were generated using AUGUSTUS v 3.3 (Stanke et al., 2008). To generate the multiple preliminary analyses necessary for AUGUSTUS, we first ran BUSCO v.4.1.1 using the endopterygota_odb10 dataset (https://busco-data.ezlab.org/v4/data/lineages/endopterygota_odb10.2020-08-05.tar.gz) on the *Agrypnia vestita* and *Hesperophylax magnus* unfiltered assemblies with with options --long -m genome and --offline. Next, we ran RepeatModeler 2.0 (Flynn et al., 2019) to identify repetitive elements. We then used RepeatMasker 4.1.0 (Smit and Hubley 2008–2015) and the output from RepeatModeler to mask all repetitive elements with hard masking turned on. We used the RepeatMasker output to create a hints file by first using the script rmOutToGFF3.pl from RepeatMasker to create a gff3 file and then using the gff2hints.pl script (https://github.com/genomecuration/JAMg/blob/master/bin/gff2hints.pl). Next, we aligned transcriptomes to each genome using BLAST-like Alignment Tool v3.6 (BLAT, Kent, 2002). We aligned the transcriptome of *P. grandis* from the 1KITE project (111126_I883_FCD0GUKACXX_L7_INShauTBBRAAPEI-22 http://www.1kite.org/) to *A. vestita* and the transcriptome from *H. occidentalis* (Wang et al., 2015) to *H. magnus*. Afterwards, we sorted the .psl file and used the script blat2hints.pl from the AUGUSTUS module. The extrinsic file contains necessary information for our sources of evidence: E (BLAT) and RM (RepeatMasker). Finally, to run AUGUSTUS we used the following input files: the assembly (hard masked), extrinsic file, merged RepeatMasker and BLAT hints files, and retrainting parameters (from BUSCO). In order to run AUGUSTUS in parallel, the script partition_EVM_inputs.pl from EVM (Haas et al., 2008) was used to create folders with one scaffold and a corresponding hints file. At this point, scaffolds that were marked as contaminated from BlobTools were removed. The output files were then concatenated in numerical order and joined into a gff file using the script join_aug_pred.pl from AUGUSTUS. Afterwards, we characterize the gene models from AUGUSTUS using ncbi-blast2.9.0+ blastp with -e-value 1e-4, -max_hsps 5, -outfmt 5, and -max_target_seqs 10. Functional annotations were assigned using Blast2GO (Götz et al., 2008).

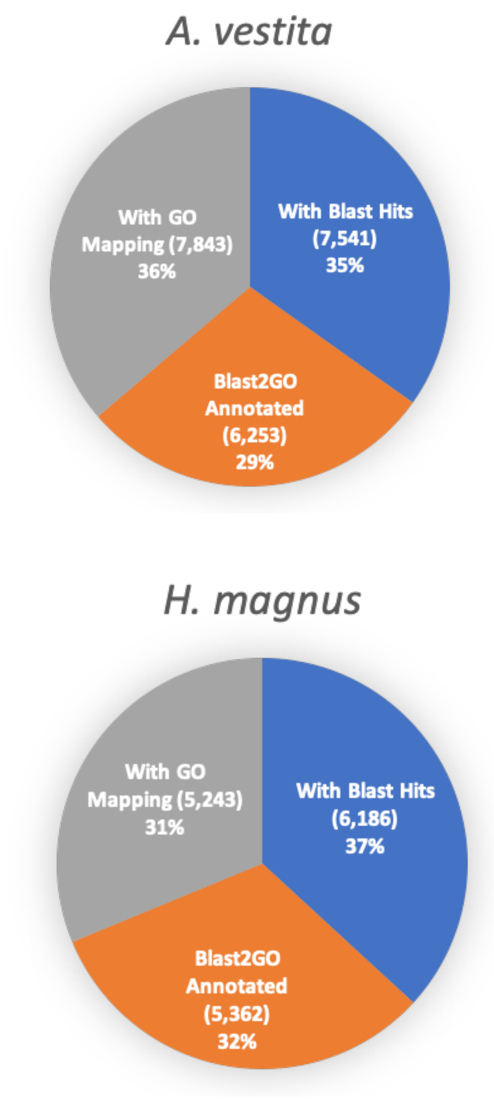


**Supplemental Figure 5:** Blast2GO Annotation Results
Pie charts showing the percentage of proteins in *A. vestita* and *H. magnus* with functional Blast2GO annotations that were verified by BLAST and mapped to GO terms compared to proteins lacking a functional annotation but verified by BLAST and mapped to GO terms or proteins only verified by BLAST.

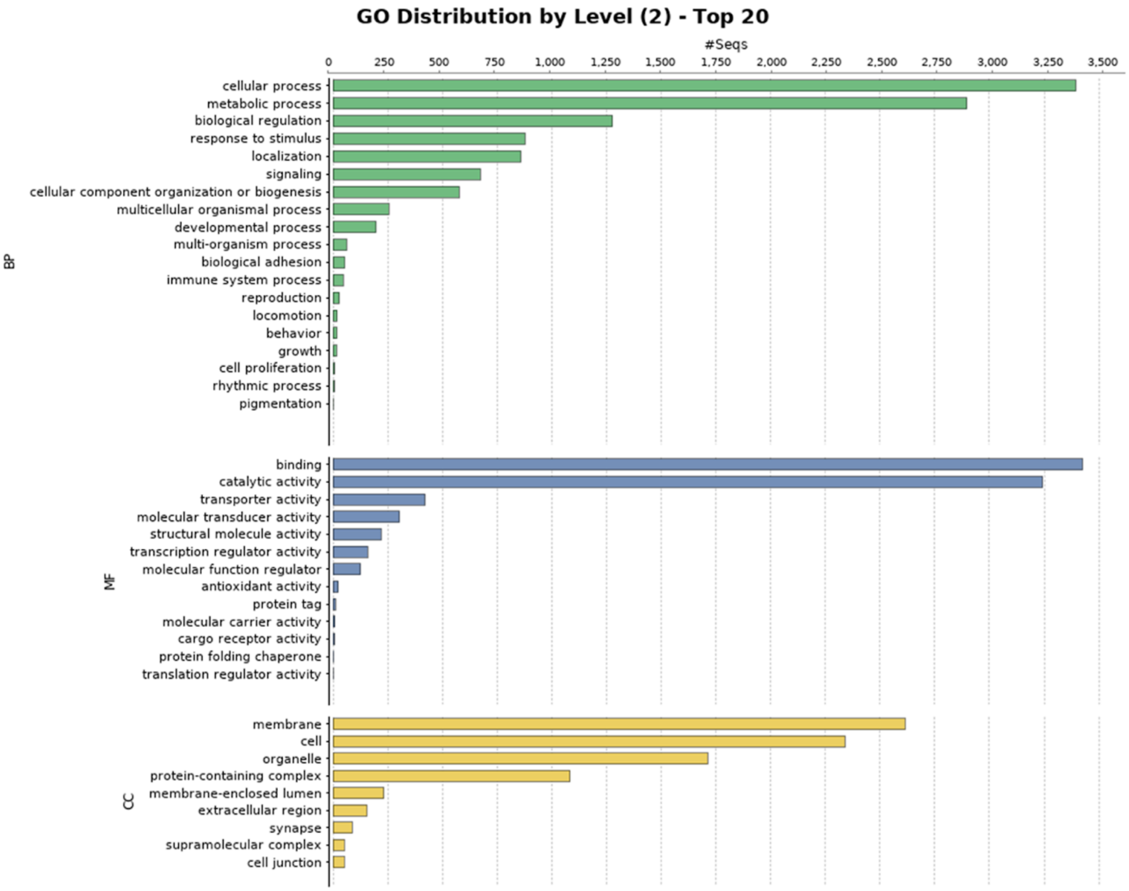


**Supplemental Figure 6:** Blast2GO Functional Annotation for *A. vestita*
Barplot showing GO terms characterized by biological process, molecular function, and cellular component. Barplots are grouped by biological process (BP), molecular function (MF), and cellular component (CC).

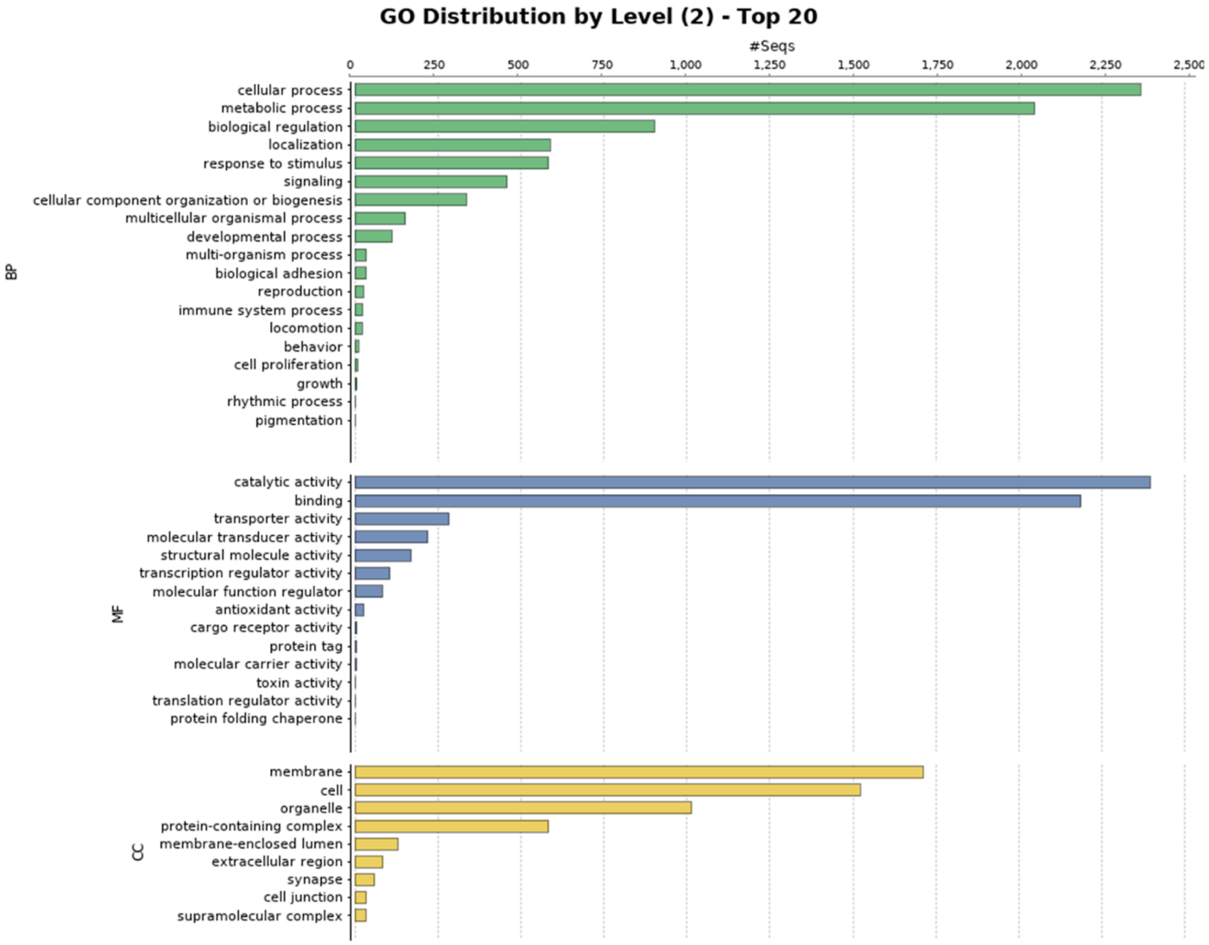


**Supplemental Figure 7:** Blast2GO Functional Annotation for *H. magnus*
Barplot showing GO terms characterized by biological process, molecular function, and cellular component. Barplots are grouped by biological process (BP), molecular function (MF), and cellular component (CC).
